# A comparison of control samples for ChIP-seq of histone modifications

**DOI:** 10.1101/007609

**Authors:** Christoffer Flensburg, Sarah A Kinkel, Andrew Keniry, Marnie E Blewitt, Alicia Oshlack

## Abstract

The advent of high-throughput sequencing has allowed genome wide profiling of histone modifications by Chromatin ImmunoPrecipitation (ChIP) followed by sequencing (ChIP-seq). In this assay the histone mark of interest is enriched through a chromatin pull-down assay using an antibody for the mark. Due to imperfect antibodies and other factors, many of the sequenced fragments do not originate from the histone mark of interest, and are referred to as background reads. Background reads are not uniformly distributed and therefore control samples are usually used to estimate the background distribution at any given genomic position. The Encyclopedia of DNA Elements (ENCODE) Consortium guidelines suggest sequencing a whole cell extract (WCE, or “input”) sample, or a mock ChIP reaction such as an IgG control, as a background sample. However, for a histone modification ChIP-seq investigation it is also possible to use a Histone H3 (H3) pull-down to map the underlying distribution of histones.

In this paper we generated data from a hematopoietic stem and progenitor cell population isolated from mouse foetal liver to compare WCE and H3 ChIP-seq as control samples. The quality of the control samples is estimated by a comparison to pull-downs of histone modifications and to expression data. We find minor differences between WCE and H3 ChIP-seq, such as coverage in mitochondria and behaviour close to transcription start sites. Where the two controls differ, the H3 pull-down is generally more similar to the ChIP-seq of histone modifications. However, the differences between H3 and WCE have a negligible impact on the quality of a standard analysis.

## 1. Introduction

As sequencing grows cheaper and more efficient (Koboldt, Steinberg, Larson, Wilson, & Mardis, 2013), ChIP-seq of histone modifications is becoming an increasingly powerful tool to understand the underlying mechanics of gene regulation (Trynka et al., 2013; Zhu et al., 2013). Studies utilizing ChIP-seq of histone modifications are now common in research, and large public data sets are also available through consortia such as ENCODE, (Consortium, 2004) or Epigenomics Roadmap (Bernstein et al., 2010). The ChIP protocol is complicated however, and it is often hard to achieve consistently high quality data (Park, 2009). A necessity of the ChIP protocol is the use of antibodies, which inherently have varying specificity (Bock et al., 2011) that can introduce unwanted features. Furthermore there are biases in the sequencing process and alignment, such as PCR amplification artefacts, GC biases and alignment artefacts. See (Park, 2009) for further discussion of error sources in ChIP-seq.

A widely accepted technique for handling these biases is the use of control samples that exhibit similar biases as the ChIP samples, but without the enrichment related to the ChIP target. The enrichment of the target histone modification can then be extracted by comparing the ChIP to the control samples. This procedure can eliminate a wide range of error sources, but hinges on the quality of the control as much as on the quality of the ChIP (Ho et al., 2011; Landt et al., 2012).

The most common control samples used are whole cell extract (WCE) or a mock pull down. The WCE is a sample of the sheared chromatin taken prior to immunoprecipitation and is often referred to as the “input”, whereas a mock pull down gives an estimation of background by pulling down an irrelevant target using a non-specific antibody such as IgG. Both of these options emulate some of the biases present in a ChIP-seq data set but a mock pull-down is believed to be more similar to the background signal of the ChIP sample as it emulates more steps in the processing of the sample. However, it can be difficult to retrieve sufficient amounts of DNA from a mock immunoprecipitation to give an accurate estimation of the background and so a WCE sample is by far the most common control sample used (Landt et al., 2012).

An alternative control for the ChIP-seq of Histone H3 modifications is an anti-H3 antibody immunoprecipitation, which closely mimics the background by enriching the sample at the location of histones (nucleosomes) along the DNA. Therefore, using an H3 pull-down as a control sample gives a measure of the enrichment in relation to the presence of a histone. For example, if a histone modification antibody had a slight affinity for all histones whether or not they had the modification, the H3 pull-down could account for this background. This is slightly different to a WCE control that not only misses the immunoprecipitation process in the protocol but also attempts to measure the density of a modified histone relative to a uniform genome. Here we investigate which background sample is the most efficient at reducing noise and extracting biologically relevant information from histone modification ChIP-seq data.

To our knowledge there are no studies comparing WCE or IgG to Histone H3 immunoprecipitation as a control sample, and there are few studies using H3 immunoprecipitation as a control. In this paper we use new unpublished data from a mouse hematopoietic stem and progenitor cell population that includes WCE, H3 and trimethylation of lysine 27 on histone H3 (H3K27me3) ChIP-seq samples, together with RNA-seq expression data, to study the differences between WCE and H3 pull down as control samples. This data set shows that H3 samples share some features with the H3K27me3 samples that are not present in the WCE sample, but that these biases do not have a significant impact in most standard analyses.

The first part of this paper studies the genome wide properties of the WCE and H3 samples and looks for differences and similarities between the two types of background samples. In the results section we firstly compare each background sample to histone modification ChIP-seq and expression data. We then compare which control is most successful in extracting correlation between histone modifications and expression.

## 2. Materials and methods

### 2.1. ChIP and cell isolation protocols

The mouse hematopoietic stem and progenitor cell population for ChIP was isolated from E14.5 foetal livers from C57BL/6 mice by fluorescence-activated cell sorting based on the following cell surface marker profile: lineage (Ter119, B220, CD5, CD3, Gr1) negative, c-Kit^+^ and Sca1^+^. Similarly, adult bone marrow hematopoietic stem and progenitor cells for RNA-seq were sorted based on lineage (Ter119, B220, CD19, Mac1, Gr1, CD2, CD3, CD8) negative, c-Kit^+^ and Sca1^+^ cell surface marker expression. Approximately 250,000 cells were used for each ChIP, whereas 30,000 to 100,000 cells were used to prepare RNA for the RNA-seq.

For chromatin immunoprecipitation, formaldehyde cross-linked cells were sonicated in a Covaris sonicator. A small fraction of sonicated material was retained as the WCE sample and the remainder was incubated with either an antibody against H3 (AbCam) or H3K27me3 (Millipore) overnight at 4°C. Immune complexes were purified by incubation with protein G beads (Life Technologies) at 4°C for an hour. Cross-links were reversed by incubation at 65°C for 4 hours before DNA fragments were purified with the ChIP Clean and Concentrator kit (Zymo). Sequencing libraries were prepared using the TruSeq DNA Sample Prep Kit (Illumina) and sequencing was performed on a HiSeq2000 (Illumina).

### 2.2. Data

We have generated ChIP-seq data for the histone mark H3K27me3 in a hematopoietic stem and progenitor cell population isolated from E14.5 mouse foetal liver from C57BL/6 mice. There are three replicates of H3K27me3 ChIP-seq, with approximately 16M, 17M and 18M reads each, two H3 ChIP-seq replicates with approximately 24M and 27M reads each, and one whole cell extract with 44M reads. We also generated three RNA-seq replicates from a hematopoietic stem and progenitor cell population isolated from the bone marrow of adult mice that had received foetal liver cell transplants with approximately 17M reads each. The reads are 100 bp single end reads sequenced by Illumina HiSeq technology. Data will be available from GEO upon publication. We used default Bowtie 2 version 2.2.3 with the --very-sensitive-local preset to align the ChIP-seq and WCE reads, and TopHat version 2.0.8 with the --b2-very-sensitive preset to align the mRNA-seq reads. Alignment was performed against mm10.

### 2.3. Methods

The aligned reads were filtered for mapping quality of 20 or more, and assigned to 100 and 1000 bp consecutive, non-overlapping bins over the genome based on the centre of the read. The smaller bin size is used in the gene coverage analysis, while the larger bin size is used for count distribution and differential enrichment analysis. For some analyses the larger library sizes are downsampled to match the smallest. We want to keep a fraction *p_i_* = *N*_min_/*N_i_* of the reads from each library *i*, where *N_i_* is the library size of library *i*, and *N*_min_ is the smal lest library size in the comparison. This is implemented by reassigning a bin with *b* reads in library *i* a new read count *b*′, randomly selected from a binomial distribution *B*(*b, p_i_*) of *b* tries with an expectation value of *p_i_b*.

For comparison purposes, we generated random counts based on a Poisson distribution (fig 1). Specifically, bins that have no reads in any sample are not assigned any reads in the random sample. The remaining B bins are assigned a random number of reads from a Poisson distribution with an expectation value of *λ* = *N*_min_/*B*.

**Figure 1.**
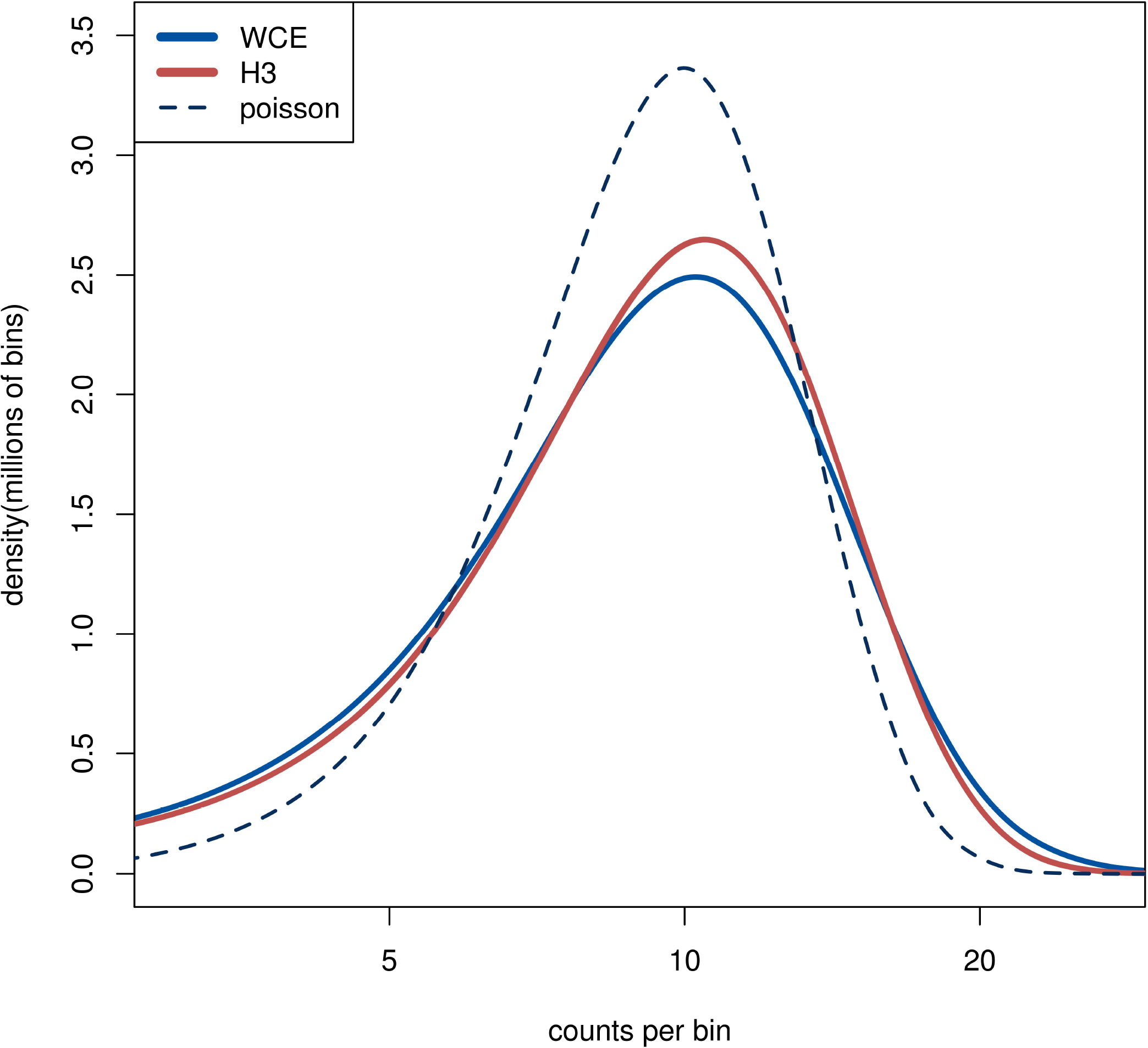
The distribution of read counts in 1kb bins over the genome. The lines from the two H3 samples are completely overlapping. The dashed line is from a random sample where the reads are distributed independently with equal probability for each bin. The curves are smoothly interpolated between the discrete data points.

The MA plots are generated using the *plotMA* function in the *limma* R package (Smyth, 2004), where the log fold change is plotted against the mean log intensity.

Differential analysis of counts between the control samples was done with limma-voom (Law, Chen, Shi, & Smyth, 2014; Smyth, 2004), with the replicated histone modification counts used in the variance estimates for each bin.

Peak finding was performed using MACS 1.4 (Zhang et al., 2008), with default parameters. Peaks from different samples were classified as overlapping if the peak regions shared at least one base pair (bp).

Expression levels were determined from read counts per million reads per kilobasepair (kb) of exon length (RPKM). The read count was increased by one read per million reads in the library (cpm increased by one). Enrichment level was determined in the same fashion, but using the full gene length (including introns) and adding 0.5 to the RPKM instead of to the cpm, to even out the background levels between the samples.

The average coverage over genes and promoters was determined from the 100 bp bin counts. Each gene was assigned 150 bins, with bin size 1/50 of the gene width, covering the gene and one gene width on either side. These bins were assigned coverages by averaging the read counts of all overlapping 100 bp bins. Bins in the same position of each gene were then averaged over genes in the same expression quartile. The average coverage in a bin was calculated as the mean over the bin and the two neighbouring bins to make the plot smoother. Genes with extremely high RPKM (larger than 100) were excluded from the analysis, to allow the mean to be calculated on a linear scale without being dominated by a few outlying genes. In our dataset, all the removed genes are ribosomal or mitochondrial.

All code for the analysis can be found in the supplementary material.

## 3. Results

Using an immunoprecipitation of Histone H3 as a background sample is attractive for accounting for uneven coverage across the genome due to both technical and biological artefacts. Specifically, an H3 pull-down not only mimics all the steps in the ChIP-seq processing but data also locates the possible regions of the genome that have the H3 protein and therefore the potential to harbour a histone modification. In order to assess the possible advantages in using an H3 control we began by comparing H3 with a standard WCE background sample.

### 3.1. Comparing background samples across the genome

As we have outlined, control samples are often used to cancel background reads that can lead to false signals in the histone modification samples. In order to verify that the controls indeed have structure beyond random sampling, such as enriched or depleted regions, we examined the distribution of reads across the genome. Initially we counted the number of reads in 1kb bins across the genome and compared the count distributions between WCE, H3 and simulated random counts from a Poisson distribution. To account for different library sizes, the larger libraries were downsampled to match the smallest one (see methods). The WCE and H3 count distributions are very similar to each other and they both display heavier tails compared with a Poisson distribution, thus implying non-random structures in the data (Figure 1). We see more bins with high counts than what is expected from a random Poisson distribution, which is an indication of enriched regions. The nearly identical distributions of the WCE and H3 samples in our data show that both control samples have similar number and size of enriched regions.

We next investigated whether these suggested non-random features coincide in location on the genome between the background samples. To do this we performed a statistical test for differential enrichment between the H3 and WCE counts in 1kb bins using limma and voom (Law et al., 2014; Smyth, 2004), and plotted the fold changes in an MA plot (Figure 2A). In the limma-voom analysis the three H3K27me3 samples are included for improved estimation of the variance between samples. The most striking difference between the WCE and H3 samples occurs at mitochondrial DNA where there is enrichment prevalent in the WCE sample but none in the H3 samples. This is expected since mitochondrial DNA is not packaged by histones and therefore is not recognised by the H3 antibody (Chen & Butow, 2005). In addition, the ribosomal gene *Rn45s* shows exceptionally high counts in both data sets, but significantly higher counts in WCE than H3. This gene is highly variable in the expression data and is likely associated with technical effects in the experiment. The remainder of the significantly different bins are intergenic repeat regions, with large counts in both WCE and H3 but modest fold changes, and are marked by RepeatMasker (Kent et al., 2002; Tempel, 2012) as repetitive elements. It is important to note that apart from the enrichment in the mitochondrial DNA we again see that the two background samples have enrichment locations that are more similar to each other than they are to a randomly generated background sample (Supplementary Figure 1). In summary the vast majority of bin counts across the genome are very similar between the control samples.

**Figure 2.**
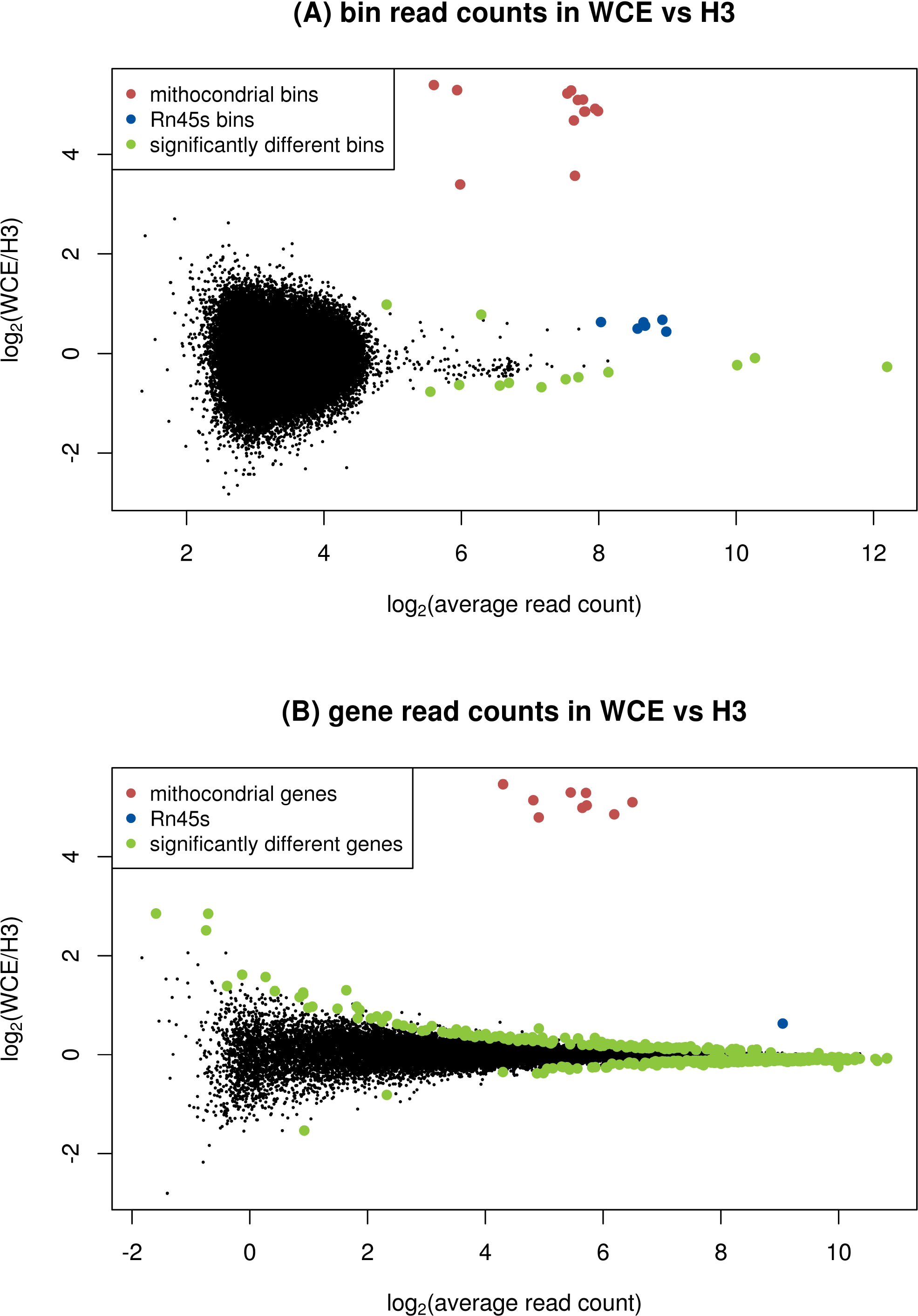
MA plots comparing enrichment between WCE and H3: (A) 1kb bins across the genome, (B) bins that span genes. Significantly different bin counts between samples are highlighted and *Rn45s* and mitochondrial bins (all significantly different) are highlighted in blue and red respectively. Significance is defined as FDR below 0.05 in a limma-voom analysis.

### 3.2. Comparing background samples over genes

We next shifted our focus to the behaviour of the background samples in and around genes, as these are often the regions of most interest in functional studies. Figure 2B shows an MA plot comparing the number of reads in each gene for WCE and H3. For this analysis significance is tested using the total number of reads across the entire gene, which for most genes gives more power than the 1kb bin count. We find 444 genes are differentially enriched with an FDR below 0.05. However, only 35 of the significantly different genes have a fold change larger than a factor 1.5. Consistent with the analysis of bin counts above, it is only the mitochondrial genes and *Rn45s* that stand out from the rest of the population as being more highly enriched in the WCE compare with the H3 sample.

### 3.3. Comparisons of detected peaks

The bin counts analysis above is not optimised to capture enrichment features that either span multiple bins or are significantly smaller than the bin size. Therefore, we next identified enriched regions in the data using MACS to call peaks in our background samples. MACS (Zhang et al., 2008) is designed to find enriched peaks in the coverage of ChIP-seq data, and scores the peaks according to significance.

MACS identified 988 peaks in the mouse WCE sample, and 872 peaks in the merged H3 samples, with 644 peaks overlapping between the samples. We found that overlapping peaks between WCE and H3 were assigned similar scores by MACS (Figure 3B, Pearson correlation 0.89). We also noted that the peaks that appeared in one of the controls, but not in the other are scored much lower than the peaks that are common between the samples (fig 3A). These two observations show that WCE and H3 have very similar peak locations.

**Figure 3.**
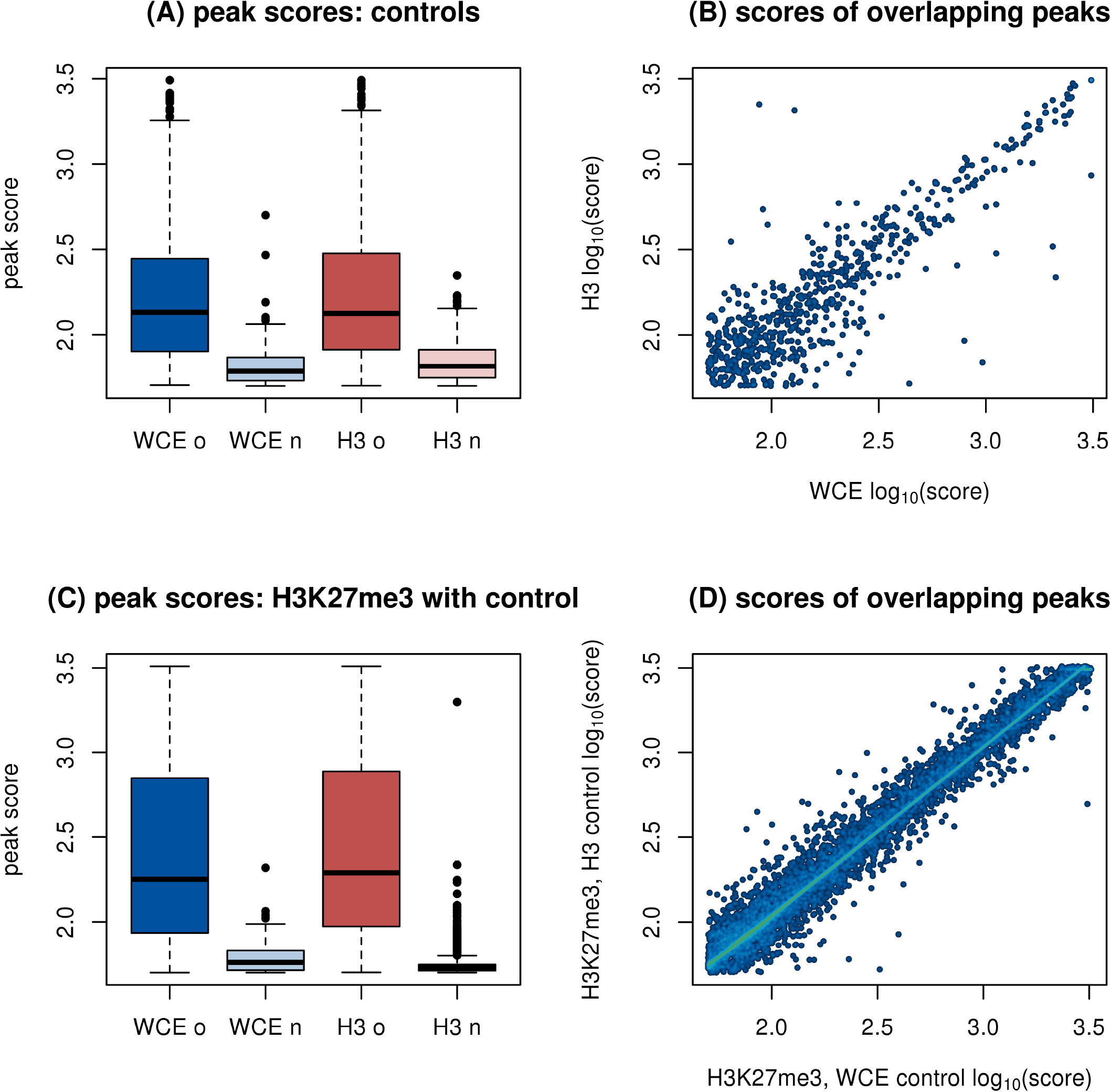
Peak scores from a MACS analysis of enriched regions. H3 and H3K27me3 replicates are merged before analysis. (A) Distributions of scores of peaks in WCE and H3. The peaks in each control are split into peaks that overlap between the two background samples (marked with “o”) or do not overlap (marked with “n”). (B) Scatter plot of scores of overlapping peaks from the WCE and H3 sample. (C) Distributions of scores of peaks in H3K27me3 with either WCE or H3 as a control. Labels on the x-axis refer to the control sample. (D) Scatter plot of scores of overlapping peaks from H3K27me3 with either WCE or H3 as control.

We next performed a standard peak finding analysis of the merged H3K27me3 samples, using either WCE or H3 as a control. For WCE and H3, MACS returned 19,683 and 21,315 regions respectively in the H3K27me3 data. The vast majority of these peaks were overlapping (19,552 regions). The enriched regions that are detected with both background samples have highly correlated peak scores (Figure 3D, Pearson correlation 0.996), and the few peaks that are unique to the analysis for only one of the background samples have very low scores (fig 3C). This shows that the small differences that exist between our WCE and H3 samples have a negligible impact in a typical peak finding analysis and would not be prioritized for follow up.

Using any of the controls on our data set improved the quality of the analysis. Without the controls, only 7493 peaks were called in the H3K27me3 samples, and many biologically relevant peaks were lost (supplementary methods and supplementary figure 2).

### 3.4. Comparison of H3K27me3 to expression with different background samples

It is known that H3K27me3 enrichment at the promoter or body of a gene is associated with low expression of that gene (Young et al., 2011). First we used expression levels estimated from our RNA-seq data to split the genes into four equal groups, or quartiles, based on expression from least to most expressed. We then plotted the average read density of genes in each quartile for each of our ChIP-seq and background samples (Figures 4A-F).

**Figure 4.**
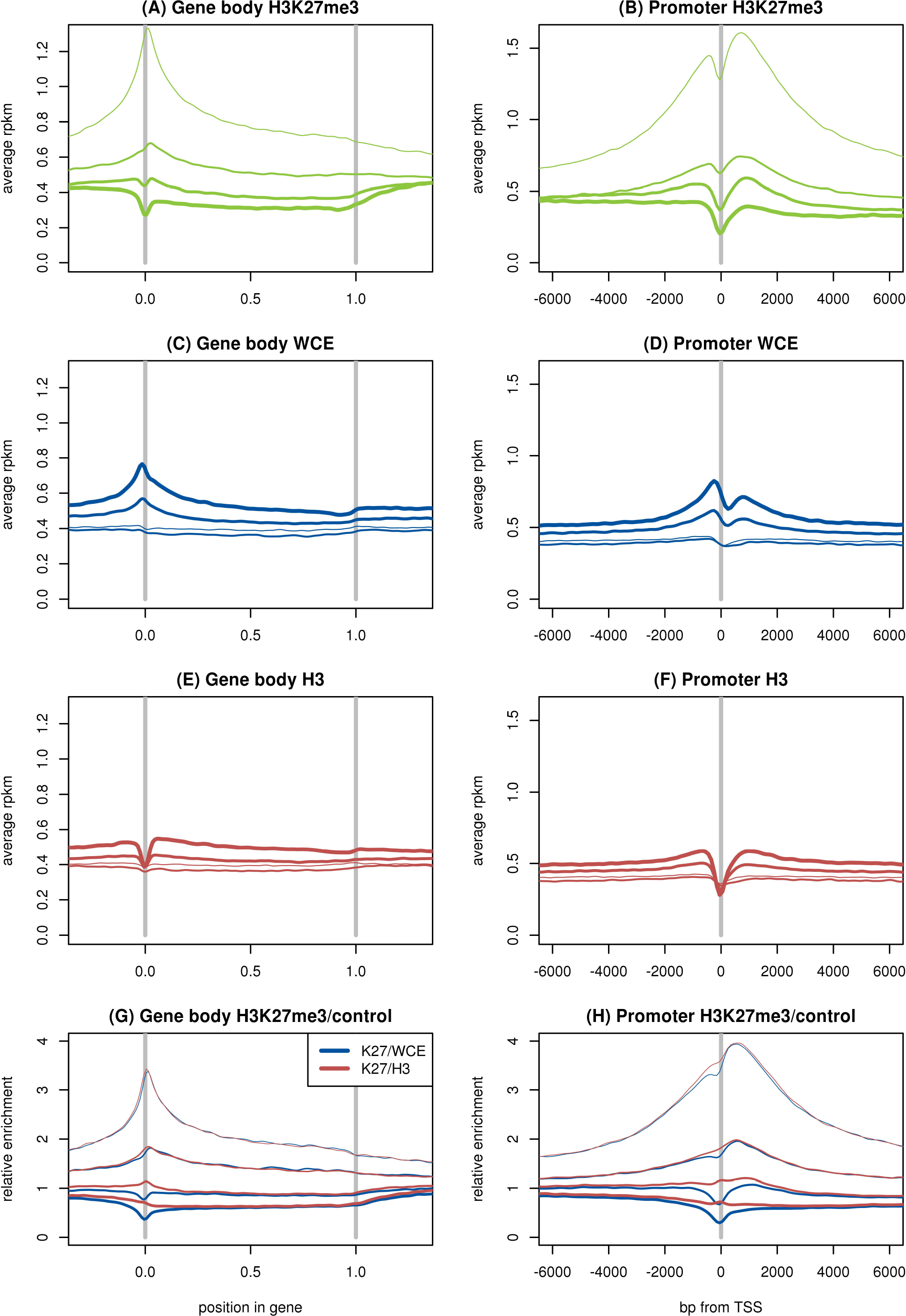
Average read density in RPKM over genes (A,C,E) and around the transcription start site (B,D,F) for H3K27me3 (A,B), WCE (C,D) and H3 (E,F). Ratio of average read densities of H3K27me3 to the background sample over genes (G) and around the TSS (H). The genes are divided up into four equal groups based on expression (RPKM), shown by thin lines for lowest expression and thicker lines for higher expression. The two H3 samples and three H3K27me3 samples are merged.

As expected, we found that H3K27me3 is more enriched across gene bodies for lowly expressed genes, and has pronounced enrichment over the transcription start site for the lower expression quartiles. By contrast, both WCE and H3 have somewhat higher coverage in highly expressed genes. The WCE sample displays a peak over the transcription start site (TSS) for the highly expressed genes, whereas the H3 and H3K27me3 samples display a dip or trough at the TSS. The dip for H3 and H3K27me3 is consistent with a depletion of nucleosomes at the TSS (Ozsolak, Song, Liu, & Fisher, 2007), which also could contribute to the peak in the WCE sample. We then divided the H3K27me3 coverage by the coverage of the controls, and found that the H3K27me3 dip was perfectly cancelled out by the H3 control, while dividing by WCE reinforced the dip (Figure 4G,H).

The perfect cancelation shows that the dips in H3K27me3 and H3 likely are from the same source. This difference between H3 and WCE is important, as the depletion of histone modifications over the TSS compared to WCE might be misinterpreted as histones around the TSS being unmodified. Using the H3 as control shows that the dip in H3K27me3 at the TSS is due to a lack of histones in general: the few histones that are there carry modifications as frequently as other histones close by. This difference between the controls is discussed further in the discussion.

One overall measure of quality of the ChIP-seq data is how well the H3K27me3 histone modification anti-correlates with gene expression. To investigate this we used the number of reads (RPKM) in the gene body or the promoter (defined as 4kb centred at the TSS) for the H3K27me3 mark (Table 1). We then calculated the same correlation after dividing by the counts (RPKM) measured in the WCE or H3 control samples. Table 1 shows the Pearson correlations between the log of the ChIP-seq enrichment at genes (RPKM or ratio of RPKMs) and log of the expression (RPKM).

**Table 1:** Pearson correlation between expression and enrichment of our H3K27me3 data and H3K27me3 and H3K36me3 from (Adli, Zhu, & Bernstein, 2010). Enrichment is measured by the logarithm of RPKM, or logarithm of ratio of RPKM in the case of the modification with control samples, and expression is measured by the logarithm of RPKM of the RNA-seq samples.

|  | WCE | H3 | H3K27me3 | H3K27me3/WCE | H3K27me3/H3 |
| --- | --- | --- | --- | --- | --- |
| Gene body | 0.3 | 0.27 | -0.23 | -0.46 | -0.44 |
| Promoter | 0.35 | 0.25 | -0.26 | -0.57 | -0.51 |
| *Adli et al.* |  |  | H3K27me3 | H3K27me3/WCE | H3K27me3/H3 |
| Gene body |  |  | -0.51 | -0.64 | -0.63 |
| Promoter |  |  | -0.49 | -0.66 | -0.63 |
| *Adli et al.* |  |  | H3K36me3 | H3K36me3/WCE | H3K36me3/H3 |
| Gene body |  |  | 0.69 | 0.59 | 0.62 |
| Promoter |  |  | 0.43 | 0.26 | 0.35 |

As expected, H3K27me3 enrichment over both the gene body and the promoter show an anti-correlation with expression. However, dividing by a control sample greatly improves the strength of the correlation for both the gene body and promoter. The WCE background improved the correlation slightly more than the H3 samples, especially for the promoter region. Both controls have a fairly strong positive correlation to expression, indicating that there likely is a positive background contribution to the H3K27me3 correlation, cancelling out part of the negative correlation. This background contribution was removed by dividing by the control samples, and the full negative correlation from the histone modification appears.

To investigate this effect further, we took data from (Adli et al., 2010) on H3K27me3 and H3K36me3 enrichment in adult mouse hematopoietic stem and progenitor cells, and calculated the correlations using our WCE and H3 as background samples. These datasets are from a cell type that is more similar to that used for our expression data, giving stronger correlations overall, however the same trends were observed in relation to the control samples. The H3K27me3 data behaved the same as in our dataset, in that both controls strengthened the anti-correlation between H3K27me3 enrichment and expression, and the WCE provided stronger anti-correlation than the H3 control. Conversely, the H3K36me3 correlation to expression was strong without controls, but weakened by the introduction of controls. Again, the effect was strongest in the promoter with WCE as control. It should be noted here that the weaker correlation in the H3K36me3 at the promoter is expected, as H3K36me3 is an elongation mark, and is normally enriched over exons rather than promoters. These results can be interpreted as both H3K27me3 and H3K36me3 having their correlation to expression influenced by the background. The control samples are both positively correlated to expression, indicating that the background likely increases the correlation for the histone modifications. This explains why the anti-correlation of expression with H3K27me3 is strengthened by the controls, while for H3K36me3 the correlation is weakened.

## 4. Discussion

We have generated ChIP-seq data for Histone H3, H3K27me3 and WCE samples from a hematopoietic stem and progenitor cell population isolated from E14.5 mouse foetal liver from C57BL/6 mice, and mRNA-seq expression data from a hematopoietic stem and progenitor cell population isolated from the bone marrow of adult mice that had received foetal liver cell transplants. We used these data, together with publicly available data, to compare WCE and H3 for their efficacy as control samples for modified histone ChIP-seq data.

Using WCE or H3 as a control yields slightly different interpretations of the corrected data. Dividing a histone modification ChIP-seq sample by a WCE background normalises the sample against genome and fragmentation biases, with no explicit regard to the underlying histones. In contrast, dividing by a H3 background estimates the enrichment relative to the position of histones in the sample. Which of these two measures is more appropriate depends on the experiment, and the type of biology being probed. However, most experiments with ChIP-seq of histone modifications do not have the precision to tell the two interpretations apart, and we will focus on which control sample can extract the most information from the experiment.

We found, as has previously been observed, that both the WCE and H3 samples have more features and structures than would be expected from randomly distributed reads. For example, both controls are more enriched over highly expressed genes than over lowly expressed genes. We compared read counts in 1kb bins between WCE and H3 and found that most features of the read distributions are similar between background samples. The notable exception to this was mitochondrial DNA where no H3 histones are present. The non-random enrichment features seen in both background samples can occur from biological sources, such as accessibility of the DNA, stability of the interaction between DNA fragments and targeted proteins as well as technical processes such as PCR artefacts and sequencing biases. Furthermore, biases can arise from the data analysis, with mappability perhaps being the most influential. These error sources are also expected to be present in ChIP-seq of histone modifications, and thus the WCE and H3 are suitable as background controls, with the majority of the structures in both WCE and H3 capable of cancelling out biases.

When the data is averaged over a large number of genes, a more relevant difference between the control samples become apparent. WCE shows more enrichment over the TSS for expressed genes, while H3 and H3K27me3 show a trough at the TSS. The trough seen in both the H3 and H3K27me3 pull-down is likely due to nucleosome depletion specifically at the TSS where the transcriptional machinery is accessing the DNA (Ozsolak et al., 2007). Possibly the peak in WCE is due to the same reason, where the WCE extraction instead will collect a larger number of reads from regions that are depleted of nucleosomes. The small dip in WCE coverage just after the TSS could then be caused by the first nucleosome, often located in roughly the same position in expressed genes as was shown in (Ozsolak et al., 2007). This difference between WCE controls and histone ChIP samples can have an impact on an analysis of expressed genes in the TSS region as can be seen in Figure 4H.

Apart from very specific analyses averaged over many genes around the TSS, the two control samples give very similar results. Using WCE or H3 as the control for peak finding in H3K27me3 ChIP-seq data returned essentially the same peaks (less than 1% of peaks were not concordant when called against WCE or H3), and the peak scores had a Pearson correlation of 0.996. We believe that in the majority of cases the choice of which background sample is used will make a negligible difference to the results. Therefore, the use of WCE or input as a standard control sample is well justified.

## Acknowledgement

This work was supported by NHMRC grants (APP105140 to AO and APP1027398 to MEB and AO). AO was supported by a Career Development Fellowship from the NHMRC. MEB was supported by an Australian Research Council, Queen Elizabeth II Fellowship. This work was made possible through Victorian State Government Operational Infrastructure Support and Australian National Health and Medical Research Council Research Institute Infrastructure Support Scheme.

